# Lipid flippases ATP9A and ATP9B form a complex and contribute to the secretory pathway from the Golgi apparatus

**DOI:** 10.1101/2024.11.13.623339

**Authors:** Tsukasa Yagi, Riki Nakabuchi, Yumeka Muranaka, Gaku Tanaka, Kazuhisa Nakayama, Hiroyuki Takatsu, Hye-Won Shin

## Abstract

Type IV P-type ATPases (P4-ATPases) serve as lipid flippases, translocating membrane lipids from the exoplasmic (or luminal) leaflet to the cytoplasmic leaflet of lipid bilayers. In mammals, these P4-ATPases are localized to distinct subcellular compartments. ATP8A1 and ATP9A, both members of the P4-ATPase family, are involved in endosome-mediated membrane trafficking, although the roles of P4-ATPases in the secretory pathway remain to be clarified. ATP9A and ATP9B are located in the *trans*-Golgi network, with ATP9A also present in endosomal compartments. This study unveiled the overlapping roles of ATP9A and ATP9B in transporting VSVG from the Golgi to the plasma membrane within the secretory pathway. Furthermore, we demonstrated that the flippase activities of ATP9A and ATP9B were crucial for transport process. Notably, we discovered the formation of homomeric and/or heteromeric complexes between ATP9A and ATP9B. The existence of the heteromeric complex notably contributed to the retention of ATP9A in the Golgi. Therefore, ATP9A and ATP9B play a role in the secretory pathway from the Golgi to the plasma membrane, forming either homomeric or heteromeric complexes.

## INTRODUCTION

Membrane traffic is facilitated by vesicular and tubular carriers, which are formed at the plasma membrane or organelles through the activities of numerous factors. These include small GTPases, coat proteins, SNARE proteins, fission proteins such as members of the dynamin family, and other accessory proteins. During the formation of these vesicular-tubular carriers and the fusion of organelles, dynamic changes in membrane shape occur. These shape changes are believed to involve alterations in the composition and distribution of lipids, as well as many of the aforementioned proteins (Graham, 2004; Graham and Kozlov, 2010; Shin et al., 2012).

P4-ATPases, a subfamily of P-type ATPases, have been implicated in the flipping of membrane lipids from the exoplasmic (luminal) leaflet to the cytoplasmic leaflet (Palmgren and Nissen, 2011). P4-ATPases are crucial for regulating the trans-bilayer lipid asymmetry of biological membranes: phosphatidylserine (PS) and phosphatidylethanolamine (PE) are primarily located in the cytoplasmic leaflet, while phosphatidylcholine (PC), sphingomyelin, and glycolipids are enriched in the exoplasmic leaflet (Devaux, 1991; Murate et al., 2015; Op den Kamp, 1979; Zachowski, 1993). Among P4-ATPases, ATP8B1, ATP8B2, and ATP10A preferentially flip PC and phosphatidylinositol (Muranaka et al., 2024; Naito et al., 2015; Takatsu et al., 2014), ATP11A and ATP11C flip PS and PE (Takada et al., 2015; Takatsu et al., 2014; Yabas et al., 2011), and ATP8A1/2 primarily flip PS (Coleman et al., 2009; Lee et al., 2015). Furthermore, ATP10B flips PC and glucosylceramide, and ATP10D specifically flips glucosylceramide (Martin et al., 2020; Roland et al., 2019). The enhanced phospholipid flipping by ATP10A at the plasma membrane promotes plasma membrane deformation and induces endocytosis (Takada et al., 2018). It is also associated with plasma membrane dynamics, including changes in cell shape (Miyano et al., 2016; Naito et al., 2015). In addition, phospholipid flipping itself, as opposed to the flipped phospholipids, is important for vesicle formation in yeast (Takeda et al., 2014). Therefore, local changes in the composition of specific phospholipids by P4-ATPases are actively involved in the process of membrane deformation.

All five yeast P4-ATPases participate in protein transport at various stages of the secretory and endocytic pathways (Muthusamy et al., 2009). Some yeast P4-ATPases interact with proteins implicated in vesicle formation, such as ADP-ribosylation factor-guanine nucleotide exchange factors (ARF-GEFs). These factors activate small GTPases, which are involved in the recruitment of coat proteins and cargo receptors to vesicles (Natarajan et al., 2009; Tsai et al., 2013; Wicky et al., 2004). P4-ATPases also play crucial roles in membrane trafficking in other organisms, including *Caenorhabditis elegans* and *Arabidopsis thaliana* (Anne-Francoise et al., 2009; Lopez-Marques et al., 2021; Poulsen et al., 2008). In mammals, ATP8A1 is necessary for the recruitment of EHD1 to endosomes and for endosome-mediated trafficking in COS-1 cells (Lee et al., 2015). ATP9A is involved in the recycling pathway from endosomes to the plasma membrane of the transferrin receptor and glucose transporter 1 (Tanaka et al., 2016). ATP9A also plays a role in the recycling of Wntless and Wnt secretion (McGough et al., 2018). Therefore, the phospholipid flipping activities of P4-ATPases are vital for membrane trafficking in eukaryotic cells.

The majority of mammalian P4-ATPases necessitate an association with CDC50 for their exit from the endoplasmic reticulum (ER) and subsequent cellular localization (Bryde et al., 2010; Coleman and Molday, 2011; Paulusma et al., 2008; Takatsu et al., 2011; van der Velden et al., 2010) {Lopez-Marques, 2014 #5982}. However, ATP9A and ATP9B, considered mammalian orthologs of yeast Neo1p, do not require interaction with CDC50 for ER-exit (Takatsu et al., 2011; Tone et al., 2020) {Saito, 2004 #2037}.

Early studies highlighted the involvement of yeast and plant P4-ATPases in the post-Golgi secretory pathway (Chen et al., 2006; Gall et al., 2002; Hua et al., 2002; Poulsen et al., 2008), but the involvement of mammalian P4-ATPases in the secretory pathway remains unknown. In this study, we examined the function of ATP9A and ATP9B, which localize to the *trans*-Golgi network (TGN), in the post-Golgi secretory pathway. We discovered that ATP9A and ATP9B redundantly contribute to the secretory pathway from the Golgi apparatus to the plasma membrane. Notably, their flippase activities appeared to be essential for protein transport. Furthermore, we demonstrated that ATP9A and ATP9B function as homomeric and/or heteromeric complexes in the transport pathway.

## RESULTS

### VSVG transport from the Golgi to the plasma membrane is delayed in ATP9A or ATP9B knockout cells

ATP9A, localized to the TGN and early/recycling endosomes, and ATP9B, localized to the TGN, prompted us to investigate their roles in post-Golgi membrane trafficking (Supplementary Figure S1A) (Takatsu et al., 2011) {Tone, 2020 #9924}. To this end, we established *ATP9A* and *ATP9B*-knockout (KO) HeLa cells using the CRISPR/Cas9 system (Tanaka et al., 2016) (Supplementary Figure S2) and transfected these KO cells with the N-terminally EGFP-tagged temperature-sensitive glycoprotein of the vesicular stomatitis virus (VSVG-tsO45) (Nishimoto-Morita et al., 2009). The cells were incubated at 40℃ overnight to accumulate the EGFP-VSVG-tsO45 (EGFP-VSVG) in the ER, then shifted to 32°C to facilitate the transport of VSVG to the plasma membrane (Figure 1A, B). To inhibit protein synthesis during the 32°C incubation, cells were treated with cycloheximide.

**Figure 1.**
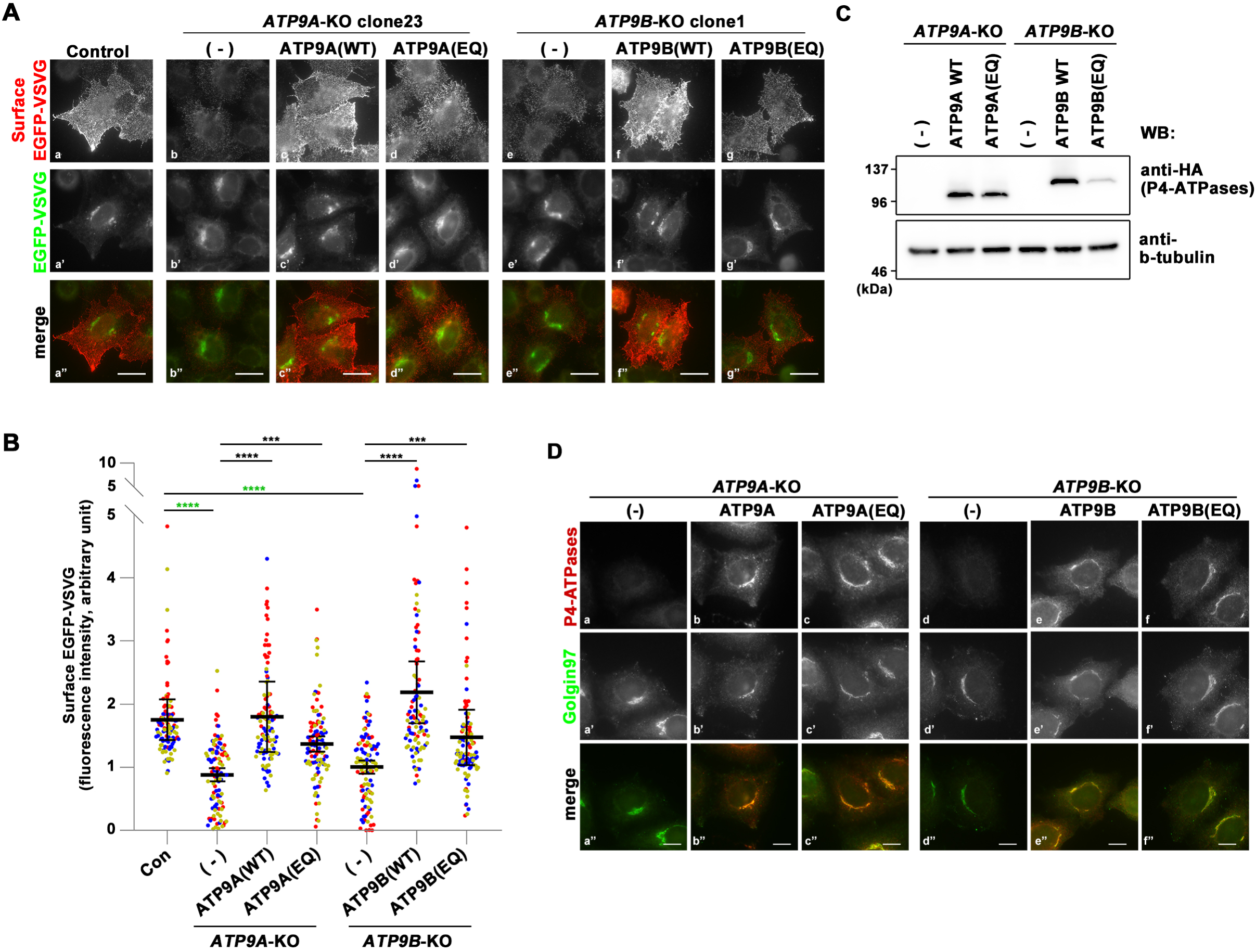
Transport of VSVG(*ts*O45) in *ATP9A*– and *ATP9B*-KO cells. Parental HeLa (Control), *ATP9A*– and *ATP9B*-KO cells, along with individual KO cells stably expressing C-terminally HA-tagged ATP9s(WT) and ATP9s(EQ) mutants, were transfected with an expression vector for N-terminally EGFP-tagged VSVG(*ts*O45). The cells were then incubated at 40°C overnight to accumulate EGFP-VSVG at the ER, followed by a shift to 32°C for 60 min to facilitate the transport of EGFP-VSVG to the plasma membrane. Cycloheximide was added during the incubation at 32°C. (**A**) To detect cell surface expression of EGFP-VSVG, cells were stained with an anti-GFP antibody under non-permeabilized conditions, followed by incubation with AlexaFluor 555-conjugated anti-rabbit secondary antibody (red). The images are representative of three independent experiments. Bars, 20 μm. (**B**) Surface fluorescence intensities (red) were quantified and normalized with EGFP fluorescence intensities (green) using ZEN software. The scatter plot displays the average ± SD from three independent experiments. Variance was assessed using a one-way ANOVA, and comparisons were made with parental HeLa cells (Control) (indicated by green asterisks) and individual KO cells (indicated by black asterisks) using Tukey’s post-hoc analysis. A total of 99 cells were quantified per sample, with each dot representing a single cell and the colors corresponding to three independent experiments. \*\*\*\**p* < 0.0001, \*\*\**p* < 0.001. The expression levels of each ATP9 protein in the individual KO cells were analyzed by immunoblotting with anti-HA and anti-β-tubulin (as an internal control) antibodies (**C**). Immunofluorescence analysis was performed with anti-HA and anti-Golgin97 antibodies, followed by incubation with Cy3-conjugated anti-rat and AlexaFluor 488-conjugated anti-mouse secondary antibodies (**D**). Bars, 10 μm.

EGFP-VSVG proteins that reached the plasma membrane were detected by incubating fixed cells with an anti-GFP antibody without permeabilization, followed by an AlexaFluor 555-conjugated secondary antibody. Thus, the red fluorescence indicates EGFP-VSVG proteins that have reached the cell surface (Figure 1Aa and a’’). When cells were incubated at 40℃ following transfection with the EGFP-VSVG construct, the EGFP signal was rarely detected, likely due to folding defects in the luminal EGFP caused by VSVG(tsO45) (Supplementary Figure 3A) (Nishimoto-Morita et al., 2009). After incubation at 32℃, the appearance of EGFP-VSVG signal in the Golgi area coincided with the emergence of red fluorescent signals, indicating its transport to the plasma membrane (Supplementary Figure 3B and C, control panels).

The red fluorescent signals were considerably reduced in *ATP9A*-KO (clone 23) and *ATP9B*-KO (clone 1) cells compared to parental control cells, suggesting a notable delay in the transport of VSVG to the plasma membrane in these KO cells (Figure 1Ab, b’’, e, and e’’). To circumvent clonal effects, we verified the delay in VSVG transport to the plasma membrane in additional clones of *ATP9A*-KO (clone 2) and *ATP9B*-KO (clone 4) cells (Supplementary Figure S3). We quantified the fluorescence intensity of EGFP-VSVG on the cell surface (red fluorescent signals), which was detected using an anti-GFP antibody. The intensity was then normalized with the total EGFP intensity to account for variations in the expression level of EGFP-VSVG. The surface VSVG signals were significantly diminished in cells depleted of ATP9A or ATP9B (Figure 1B). To validate the specific effects of ATP9A and ATP9B, we established individual KO cells stably expressing C-terminally HA-tagged ATP9A(WT) or ATP9B(WT). The delay in VSVG transport to the plasma membrane was rescued by the expression of ATP9A(WT) and ATP9B(WT) in *ATP9A*-KO and *ATP9B*-KO cells, respectively (Figure 1Ac, c’’, f, and f’’ and B). The results suggest that ATP9A and ATP9B play a role in the efficient transport of VSVG to the plasma membrane. Subsequently, we investigated whether the flippase activities of ATP9A and ATP9B are required for VSVG transport to the plasma membrane. To address this, we established individual KO cells stably expressing HA-tagged ATP9A(E195Q) or ATP9B(E272Q) mutants, which are ATPase-deficient mutants (Anthonisen et al., 2006; Palmgren and Nissen, 2011; Takatsu et al., 2014). The transport of VSVG to the plasma membrane was restored by the expression of the ATPase-deficient mutant, albeit to a slightly lesser extent compared to cells expressing ATP9s(WT) (Figure 1Ad, d’’, g, and g’’ and B). Therefore, a straightforward interpretation could be that the flippase activities of ATP9A and ATP9B are not essential for VSVG transport to the plasma membrane. We confirmed the expression of ATP9A-HA, ATP9A(E195Q)-HA, ATP9B-HA, and ATP9B(E272Q)-HA through immunoblot and immunofluorescence analyses (Figure 1C and Db, c, e, and f).

### The VSVG transport requires the flippase activity of ATP9A and ATP9B

Apart from the possibility that the flippase activities of ATP9A and ATP9B are not essential for VSVG transport, we investigated another potential scenario; ATP9A and ATP9B might work cooperatively by forming a complex. We hypothesized that the residual endogenous ATP9B in *ATP9A*-KO cells could interact with the exogenous ATP9A(E195Q) mutant, partially mitigating the delay in VSVG transport to the plasma membrane. To test this hypothesis, we generated *ATP9A* and *ATP9B* double KO (DKO) cells by depleting *ATP9B* in *ATP9A*-KO cells using the CRISPR/Cas9 system (Supplementary Figure S4). Subsequently, we established DKO cells stably expressing ATP9A(WT), ATP9B(WT), ATP9A(E195Q), and ATP9B(E272Q). As anticipated, the transport of VSVG to the plasma membrane was delayed in the DKO cells (Figure 2Ab and b’’). We quantified the fluorescence intensity of EGFP-VSVG on the cell surface (red fluorescent signals), which was detected with an anti-GFP antibody, and normalized it with the total EGFP intensity to account for variations in EGFP-VSVG expression levels (Figure 2B). The transport of VSVG to the plasma membrane was restored by the expression of ATP9A(WT) or ATP9B(WT) in the DKO cells (Figure 2A c, c’’, e, and e’’ and B), suggesting the redundant function of ATP9A and ATP9B. Importantly, the VSVG transport was not restored by the expression of ATP9A(E195Q) or ATP9B(E272Q) mutants in the DKO cells (Figure 2Ad, d’’, f, and f’’ and B), even though the E-to-Q (EQ) mutants could restore the transport in the single KO cells (Figure 1A and B).

**Figure 2.**
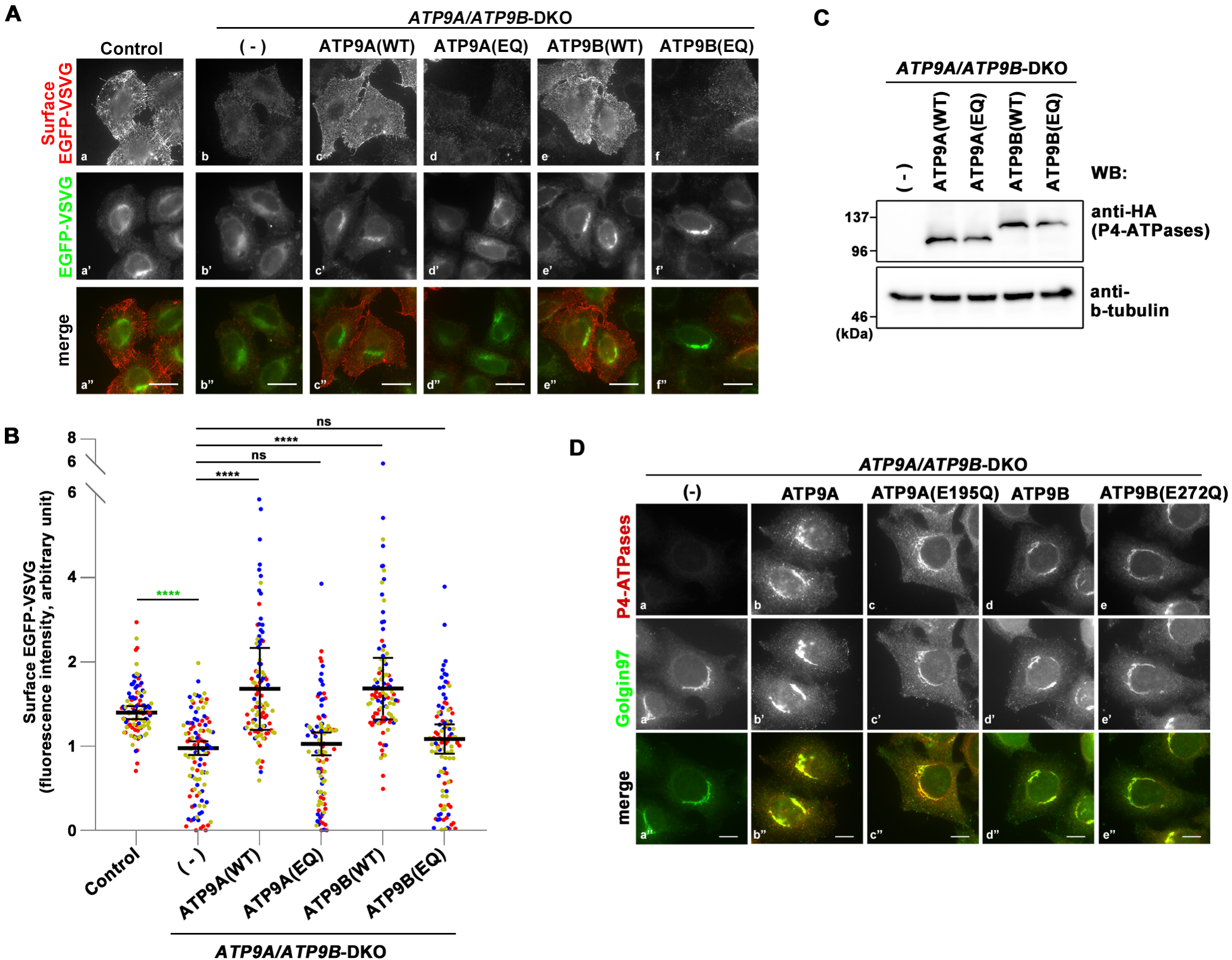
Transport of VSVG(*ts*O45) in *ATP9A/ATP9B*-DKO cells. Parental HeLa (Control), *ATP9A/ATP9B*-DKO cells, and DKO cells stably expressing C-terminally HA-tagged ATP9s(WT) and ATP9s(EQ) mutants were transfected with an expression vector for N-terminally EGFP-tagged VSVG(*ts*O45). The cells were then incubated at 40°C overnight and shifted to 32°C for 60 min to allow the transport of EGFP-VSVG to the plasma membrane. Cycloheximide was added during the incubation at 32°C. (**A**) To detect cell surface expression of EGFP-VSVG, cells were stained with an anti-GFP antibody under non permeabilized conditions, followed by incubation with AlexaFluor 555-conjugated anti-rabbit secondary antibody (red). The images are representative of three independent experiments. (**B**) Surface fluorescence intensities (red) were quantified and normalized with EGFP fluorescence intensities (green) using ZEN software. The scatter plot displays the average ± SD from three independent experiments. Variance was assessed using a one-way ANOVA, and comparisons were made with parental HeLa cells (Control) (indicated by green asterisks) and DKO cells (indicated by black asterisks) using Tukey’s post-hoc analysis. A total of 99 cells were quantified per sample, with each dot representing a single cell, and colors corresponding to three independent experiments. \*\*\*\**p* < 0.0001, ns, not significant. The expression levels of each ATP9 protein in the DKO cells were analyzed by immunoblotting with anti-HA and anti-β-tubulin (as an internal control) antibodies (**C**). Immunofluorescence analysis was performed with anti-HA and anti-Golgin97 antibodies, followed by incubation with Cy3-conjugated anti-rat and AlexaFluor 488-conjugated anti-mouse secondary antibodies (**D**). Bars, 10 μm.

We also verified the ER-to-Golgi transport of VSVG using C-terminally EGFP-tagged VSVG (Supplementary Figure S5), as N-terminally EGFP-tagged VSVG showed minimal EGFP signals in the ER upon incubation at 40°C due to folding issues (Supplementary Figure S3A). Given that protein export from the TGN is inhibited at 19.5℃ (Griffiths and Simons, 1986), we conducted a time course experiment at this temperature after 40°C incubation. The appearance of VSVG-EGFP in the Golgi apparatus was similar in control and DKO cells, indicating that ER-to-Golgi transport was unaffected in DKO cells (Supplementary Figure S5). The expression levels of these ATP9 proteins were validated through immunofluorescence and immunoblot analyses (Figure 2C and D).

These findings strongly imply that 1) the flippase activity of ATP9A or ATP9B is crucial for VSVG transport from the Golgi to the plasma membrane, and 2) ATP9A and ATP9B might function in VSVG transport as complexes.

### ATP9A and ATP9B form either homomeric or heteromeric complexes

To investigate the complex formation between ATP9A and ATP9B, we established cells that stably coexpress HA-tagged and FLAG-tagged ATP9A and ATP9B, including their EQ mutants, in various combinations (Figure 3A). We also established cells stably expressing HA-tagged ATP9A or ATP9B, along with FLAG-tagged ATP11C, a P4-ATPase family member, as a negative control. We then performed coimmunoprecipitation analysis on the cell lysates. As depicted in Figure 3A, FLAG-tagged ATP9A, ATP9B, and their EQ mutants were coimmunoprecipitated with ATP9A-HA in cells coexpressing ATP9A-HA and various FLAG-tagged P4-ATPases (lanes 1–4). A similar pattern was observed with ATP9B-HA (lanes 6–9). However, FLAG-tagged ATP11C was not coimmunoprecipitated with either ATP9A-HA or ATP9B-HA (lanes 5 and 10). These findings suggest that ATP9A and ATP9B form specific homomeric and heteromeric complexes. Moreover, ATPase deficient mutants (EQ) of ATP9A and ATP9B can also form complexes with ATP9A (lanes 2 and 7) and ATP9B (lanes 4 and 9) (Figure 3A). As previously hypothesized, the residual endogenous ATP9B in *ATP9A-*single KO cells and ATP9A in *ATP9B-*single KO cells can form heteromeric complexes with the exogenously introduced EQ mutants of ATP9A and ATP9B, respectively.

**Figure 3.**
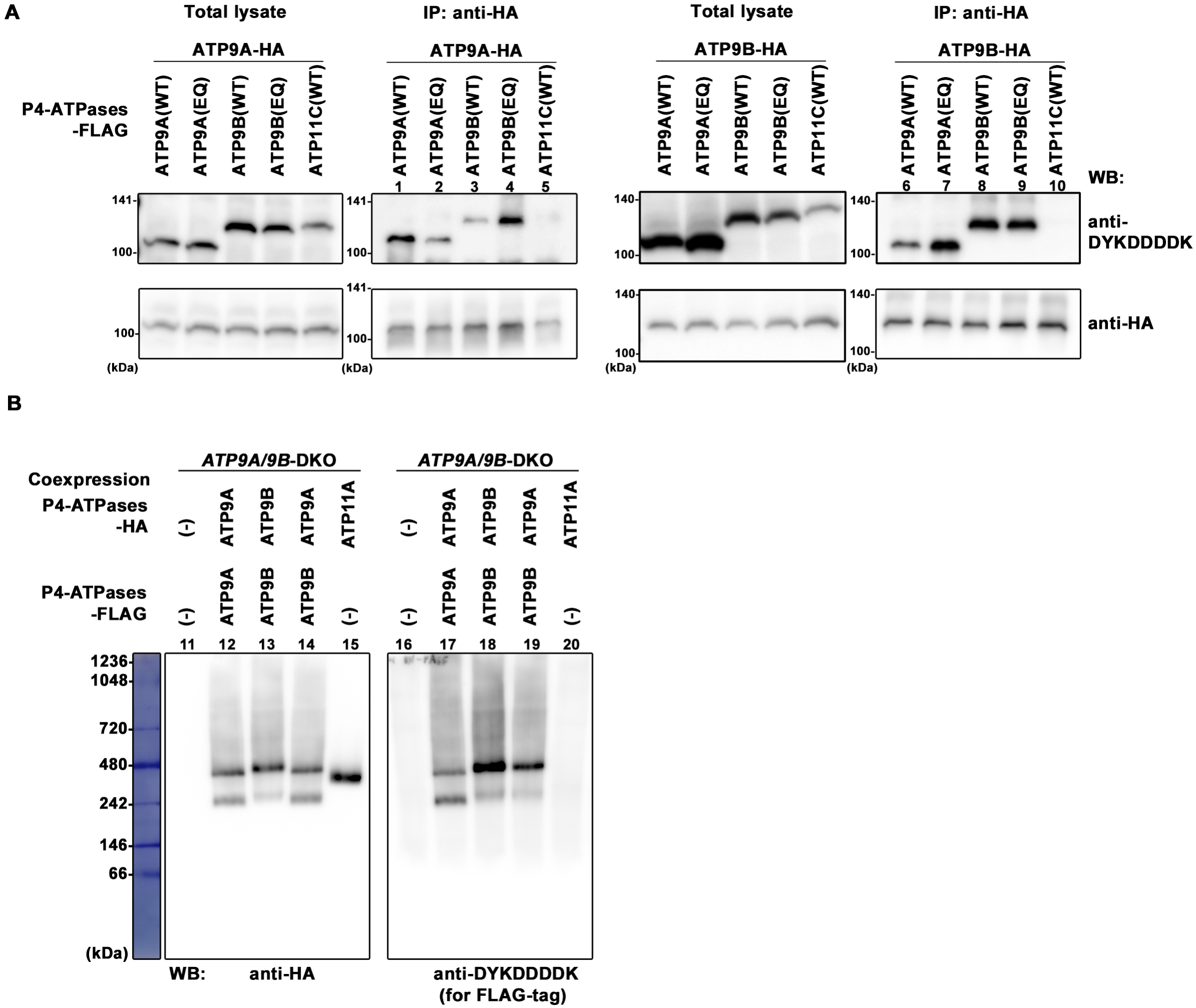
Formation of a homomeric and/or heteromeric complex between ATP9A and ATP9B. (**A**) HeLa cells, stably coexpressing HA-tagged ATP9A or ATP9B along with FLAG-tagged ATP9s, their EQ mutants, or ATP11C as indicated, were lysed. Immunoprecipitation was performed using an anti-HA antibody. Subsequently, total lysates (10% of input) and bound materials were subjected to SDS-PAGE and immunoblotting using anti-HA and anti-DYKDDDDK (for FLAG tag) antibodies. (**B**) ATP9A/ATP9B-DKO cells, stably coexpressing HA-tagged ATP9s and FLAG-tagged ATP9s in combinations as indicated, or expressing HA-tagged ATP11A alone, were lysed and subjected to BN-PAGE. Immunoblotting was performed using anti-HA and anti-DYKDDDDK antibodies.

Next, we explored the interaction between ATP9A and ATP9B under nondenaturing conditions using blue native PAGE (BN-PAGE) (Figure 3B). To avoid interference from endogenous ATP9A and ATP9B, we utilized *ATP9A/9B* DKO cells stably coexpressing HA– and FLAG-tagged ATP9A, HA– and FLAG-tagged ATP9B, or HA-tagged ATP9A and FLAG-tagged ATP9B. We also established *ATP9A/9B* DKO cells stably expressing ATP11A-HA as a control. After BN-PAGE of the cell lysates, we conducted immunoblot analysis with antibodies for the HA– and FLAG-tags (Figure 3B). ATP9A-HA and ATP9B-HA showed migration at approximately 470∼480 kDa and 240 kDa, respectively, with ATP9B-HA slightly higher than ATP9A-HA (lanes 12 and 13). ATP9A-FLAG and ATP9B-FLAG showed similar migration patterns to their HA-tagged counterparts (lanes 17 and 18), suggesting potential homomeric complex formation. ATP11A-HA exhibited migration at approximately 400 kDa, indicating its heteromeric complex formation with CDC50A (lane 15) (Takatsu et al., 2011), similar to ATP11C {Miyata, 2022 #11748}. ATP9A and ATP9B do not interact with CDC50A (Takatsu et al., 2011; Tone et al., 2020). Moreover, in cells coexpressing ATP9A-HA and ATP9B-FLAG, both ATP9 proteins appeared to be of similar size (lanes 14 and 19), although the upper band of ATP9B-FLAG showed greater intensity than its lower band (lane 19). These findings strongly support our hypothesis that ATP9A and ATP9B form either homomeric or heteromeric complexes.

### ATP9B contributes to the localization of ATP9A to the Golgi complex

During the experiment, we unexpectedly noted changes in the cellular localization of ATP9A, which was stably expressed in *ATP9B*-KO and DKO cells, compared to control cells (Figure 4A). Specifically, ATP9A-positive endosomal puncta significantly increased in *ATP9B*-KO and DKO cells (Figure 4Ac–f), while no significant change was observed in *ATP9A*-KO cells (Figure 4Ab) compared to parental HeLa cells (Figure 4Aa). The expression level of ATP9A was comparable across these cells (Figure 4B). We quantified the peripheral ATP9A-positive endosomal area and fluorescence intensity using ZEN software (Figure 4C and D). For this, we subtracted the fluorescence signal of ATP9A in the Golgi by excluding the ROI corresponding to the Golgin97-positive area (Figure 4Aa’–d’). The area and fluorescence intensity of ATP9A-positive endosomes showed a significant increase in cells lacking ATP9B (*ATP9B*-KO and DKO) (Figure 4C and D). This suggests that ATP9B contributes to either the retention or retrieval of ATP9A at the TGN, further supporting the formation of a heteromeric complex between ATP9A and ATP9B, as shown in Figure 3. Peripheral ATP9A is partially colocalized with EEA1 and transferrin receptor (TfnR), markers of early and recycling endosomes (Figure 4Ae’’, e’’’, f’’, and f’’’).

**Figure 4.**
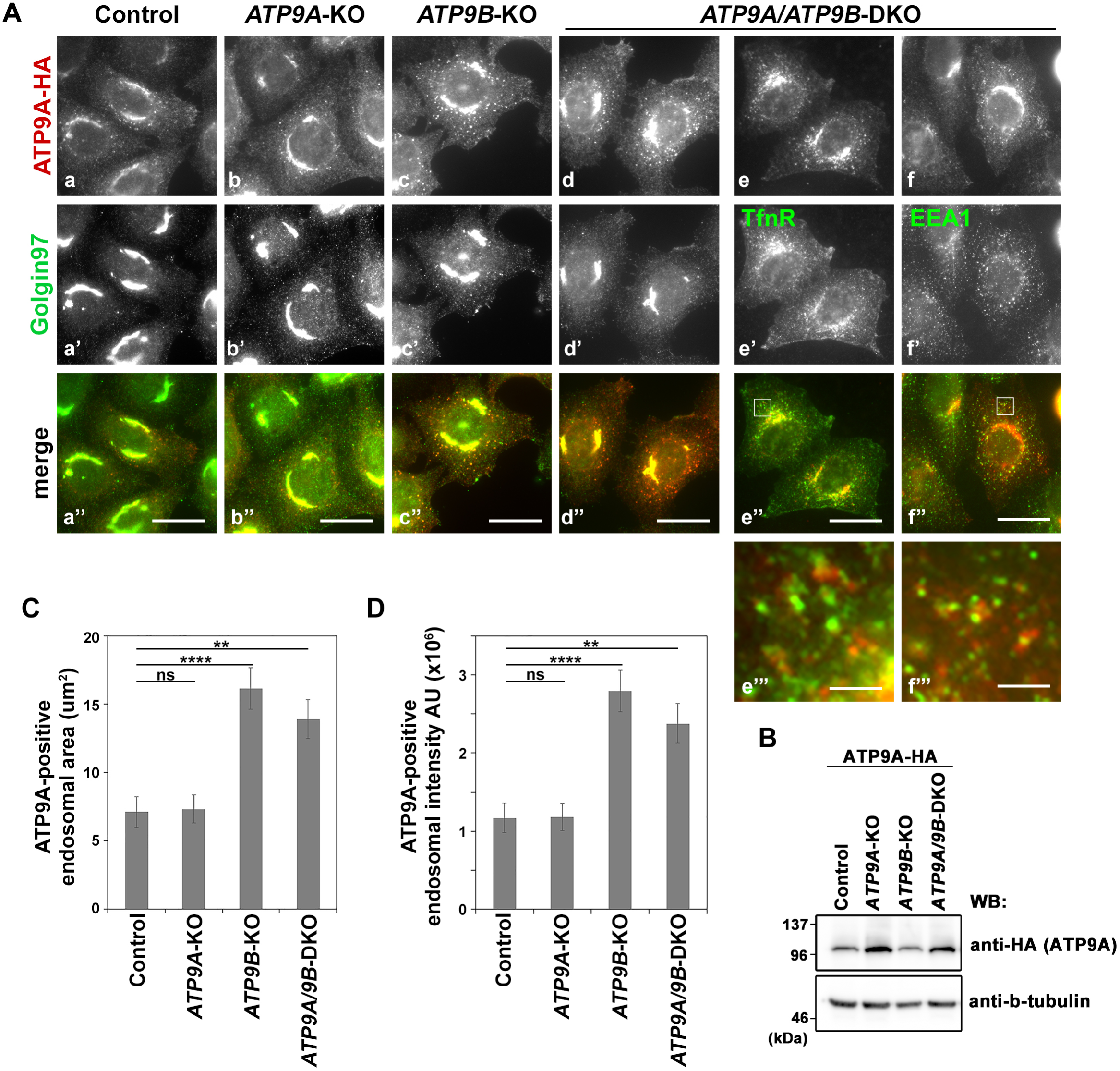
Changes in ATP9A endosomal localization in cells lacking ATP9B. (**A**) Parental HeLa (Control), *ATP9A*-KO, *ATP9B*-KO, and *ATP9A/ATP9B*-DKO cells, stably expressing HA-tagged ATP9A, were fixed, permeabilized, and then incubated with anti-HA and anti-Golgin97 antibodies (a marker for the Golgi apparatus), followed by Cy3-conjugated anti-rat and AlexFlour 488-conjugated anti-mouse secondary antibodies. Bars, 20 µm. (**B**) The expression levels of ATP9A-HA in individual cells were analyzed by immunoblotting with anti-HA and anti-β-tubulin (as an internal control) antibodies. The ATP9A-positive peripheral endosomal area (**C**) and endosomal fluorescence intensity (**D**) were quantified using ZEN software. To exclude the ATP9A-positive perinuclear Golgi area, the ROI corresponding to the Golgin97 area was subtracted. The graph represents data from two independent experiments and displays the average ± SE of over 30 cells in each sample. Variance was assessed using a one-way ANOVA, and comparisons were made with parental HeLa cells (control) using Tukey’s post-hoc analysis. \*\*\*\**p* < 0.0001, \*\**p* < 0.01, ns, not significant.

Considering the observed changes in localization of ATP9A in cells lacking ATP9B, we examined other organelle markers, including Golgi markers. TGN46 and SNARE proteins, Vti1a and syntaxin 16, cycle between the TGN and the plasma membrane via endosomes. γ-adaptin, localizing to the TGN and endosomes, facilitating trafficking between them. EEA1 and Lamp-1 serve as markers for early and late endosomes, respectively. None of these markers showed significant changes in *ATP9B*-KO, DKO, and *ATP9A*-KO cells compared to control cells (Figure 5). Thus, it seems that ATP9A and ATP9B are not essential for the recycling of at least TGN46 and some SNARE proteins between the TGN and the plasma membrane, or for the recruitment of TGN-localizing proteins like AP-1, Arl1, and arfaptins (Figure 5 and Supplementary Figure S6).

**Figure 5.**
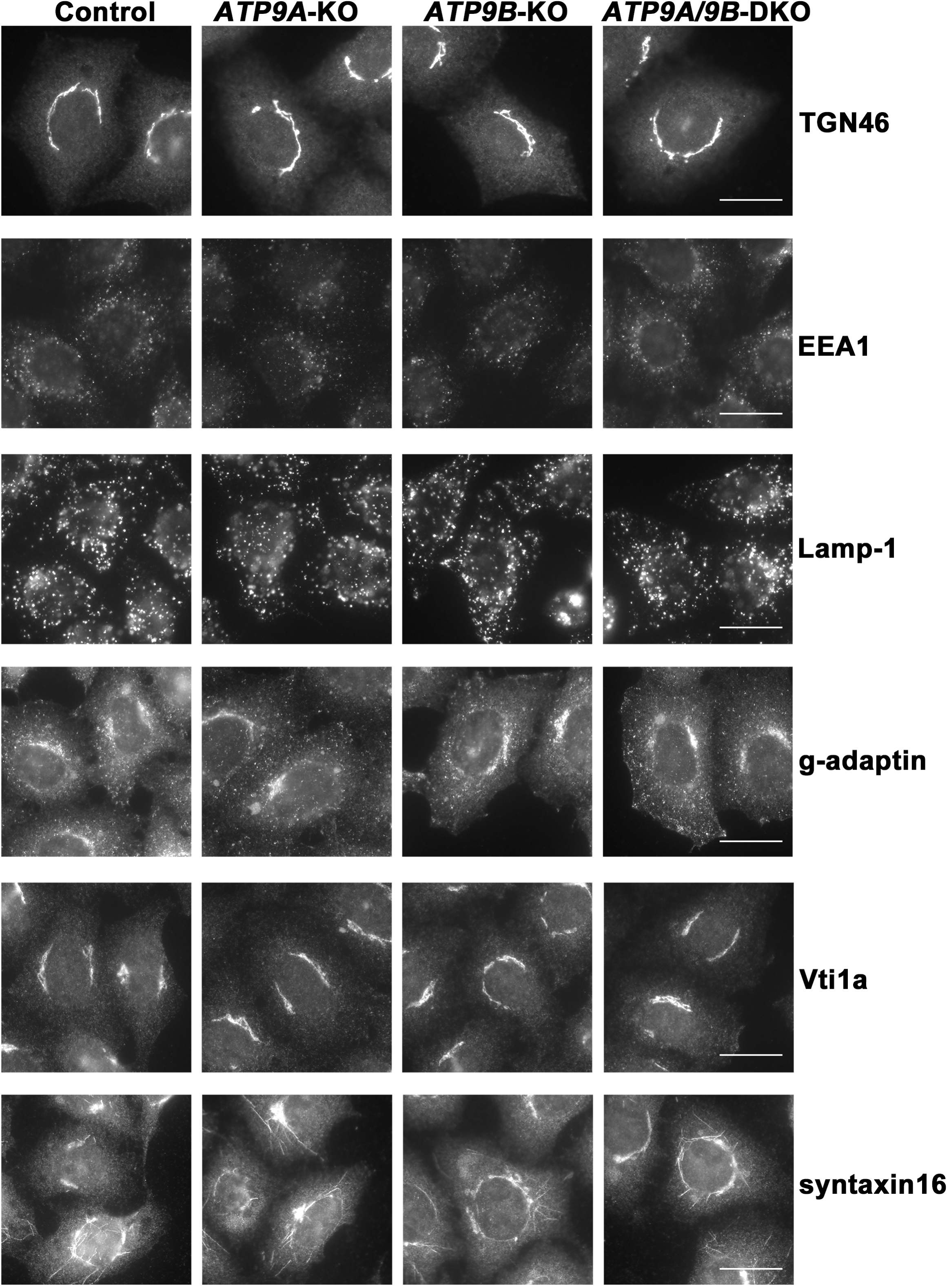
Unaltered distribution of membrane traffic–related proteins in cells depleted of ATP9A and/or ATP9B. Parental HeLa (Control), *ATP9A*-KO, *ATP9B*-KO, and *ATP9A/ATP9B*-DKO cells were fixed, permeabilized, and incubated with antibodies against TGN46, EEA1, Lamp-1, γ-adaptin, Vti1a, or syntaxin16. Bars, 20 µm.

## DISCUSSION

In this study, we demonstrated that ATP9A and ATP9B, localized to the TGN, are essential for the efficient transport of VSVG from the Golgi to the plasma membrane. Despite the integrity of cellular compartments, including the Golgi complex, remaining intact in the single KO and DKO cells, VSVG transport was delayed in the KO cells. Moreover, 1) the delay in VSVG transport was observed in both ATP9A-KO and ATP9B-KO cells, with no significantly additive effects in DKO cells. 2) The expression of either ATP9A or ATP9B in the DKO cells successfully restored the delayed VSVG transport. These findings suggest a redundant function of ATP9A and ATP9B in facilitating VSVG transport from the Golgi to the plasma membrane. We previously demonstrated that enhanced flipping activity at the plasma membrane, induced by ATP10A expression, increases membrane curvature and promotes integrin endocytosis (Naito et al., 2015; Takada et al., 2018). In yeast, lipid flipping activity is necessary for vesicle transport (Takeda et al., 2014). Thus, the flipping activities of ATP9A and ATP9B likely contribute to membrane deformation at the TGN, facilitating efficient vesicle formation at the TGN. Interestingly, we observed that the expression of ATP9A(EQ) and ATP9B(EQ) mutants partially rescued the delayed VSVG transport in single KO cells but not in DKO cells. This observation leads to two key insights: First, the flippase activity of either ATP9A or ATP9B is crucial for efficient VSVG transport. Second, ATP9A and ATP9B likely operate as a heteromeric complex. Indeed, ATP9A and ATP9B can form homomeric and heteromeric complexes. While either ATP9A or ATP9B can form a homomeric complex in *ATP9B*-KO or *ATP9A*-KO cells, respectively, VSVG transport to the plasma membrane was delayed in these KO cells. This suggests that the heteromeric complex may be more efficient in facilitating VSVG transport. Although the mechanism by which the expression of either ATP9A or ATP9B rescues the delay in VSVG transport in DKO cells remains to be elucidated, overexpression of either ATP9A or ATP9B could potentially lead to restoration.

Most mammalian P4-ATPases form a heteromeric complex with CDC50A, a chaperone-like protein, with the exception of ATP9A and ATP9B (Takatsu et al., 2011). Interaction with CDC50A is necessary for exit from the ER and localization to cellular compartments and the plasma membrane (Bryde et al., 2010; Takatsu et al., 2011; van der Velden et al., 2010). However, ATP9A and ATP9B do not require interaction with CDC50A for their localization to the TGN or endosomes. Consequently, it is plausible to speculate that ATP9A and ATP9B form a homomeric and heteromeric complex rather than interacting with CDC50A.

While the endosomal localization of ATP9A increased in cells lacking ATP9B, the distribution of Golgin97, a TGN marker, remained consistent. Additionally, the distribution of other proteins related to membrane trafficking, such as SNAREs and clathrin adaptor proteins, was not significantly altered in cells depleted of ATP9A and ATP9B. Therefore, ATP9B likely contributes to the retention and/or retrieval of ATP9A to the TGN, possibly through interaction with ATP9A.

ADP-ribosylation factors (ARFs) are central to vesicular transport from the Golgi apparatus. The ARF-like protein ARL1, which localizes to the TGN, plays a crucial role in Golgi-mediated membrane trafficking through the recruitment of its effectors, Arfaptin 1 and 2 (Man et al., 2011). Arfaptins possess the BAR domain, which senses membrane curvature and promotes membrane tubulation (Nakamura et al., 2012). ARL1 is essential for the secretory pathway in Drosophila (Torres et al., 2014), and arfaptins are involved in the secretory pathway, particularly in secretory granule biogenesis (Cruz-Garcia et al., 2013; Gehart et al., 2012). Moreover, yeast Arl1p interacts with Drs2p, a flippase localizing to the Golgi apparatus (Graham, 2013; Hsu et al., 2014; Tsai et al., 2013), and exhibits genetic and biochemical links to Neo1p, an ATP9 ortholog in yeast (Wicky et al., 2004). Therefore, ATP9A/ATP9B, ARL1, and arfaptins may be functionally linked in the secretory pathway from the TGN. However, the TGN localization of Arl1 and arfaptins is not affected by the depletion of either ATP9A, ATP9B, or both.

## MATERIALS AND METHODS

### Plasmids

Expression vectors encoding ATP9A and ATP9B, tagged at the C-terminus with HA-or FLAG, and their E-to-Q mutants were constructed as previously described (Takatsu et al., 2011; Tone et al., 2020). Each cDNA was transferred to pMXs-neo-DEST-HA, pMXs-blast-DEST-HA, and pMXs-puro-DEST-FLAG expression vectors using the Gateway system (Invitrogen). The pMXs-neo-vector and pEF-gag-pol plasmid were kindly provided by Toshio Kitamura (University of Tokyo). The pCMV-VSVG-Rsv-Rev plasmid (RDB04393) was a generous gift from Hiroyuki Miyoshi (RIKEN BRC). The pCAG-based vector for ATP9A expression with a C-terminal HA tag was previously described (Takatsu et al., 2011). Expression vectors for N-terminally EGFP-tagged VSVG (*ts*O45) (pCIpreEGFP-VSVG) (Nishimoto-Morita et al., 2009) and C-terminally EGFP-tagged pcDNA3-VSVG-EGFP, provided by Jennifer Lippincott-Schwartz (National Institutes of Health) (Presley et al., 1997), were transfected into HeLa cells using FuGene6 (Promega).

### Antibodies and reagents

The antibodies used in this study were sourced as follows: monoclonal mouse anti-EEA1 (clone 14), anti-Lamp-1 (H4A3), and anti-Golgin97 (CDF4) from BD Biosciences; monoclonal mouse anti-γ-adaptin (100.3) from Sigma; monoclonal mouse anti-Arfaptin2 (2B5) from Abnova; monoclonal mouse anti-DYKDDDDK from Wako; monoclonal rat anti-HA (3F10) from Roche Applied Science; polyclonal rabbit anti-Vti1a and anti-syntaxin16, from SySy; polyclonal rabbit anti-GFP from Molecular Probes; polyclonal goat anti-Arfaptin1 (I-19) from SantaCruz; Alexa Fluor–conjugated secondary antibodies from Molecular Probes; Cy3-and horseradish peroxidase–conjugated secondary antibodies from Jackson ImmunoResearch Laboratories. The polyclonal rabbit anti-Arl1 was previously described (Shin et al., 2005). The polyclonal rabbit anti-TGN46 (Kain et al., 1998) was a generous gift from Minoru Fukuda (Sanford-Burnham Medical Research Institute).

### Cell Culture, transfection, establishment of stable cells, and immunofluorescence

HeLa cells were cultured in a minimal essential medium supplemented with 10% heat-inactivated fetal bovine serum. Parental HeLa cells and KO cells stably expressing P4-ATPases or their EQ mutants, tagged at the C-terminus with HA or FLAG, were established via retroviral infection as previously described (Takatsu et al., 2014). The infected cells were selected in a medium containing G418 (1 mg/mL), puromycin (1 μg/mL), or blasticidin (15 μg/mL). The transport of EGFP-tagged VSVG to the cell surface or to the Golgi was examined as previously detailed (Nishimoto-Morita et al., 2009; Shin et al., 2005). In brief, cells transfected with pCIpreEGFP-VSVG or pcDNA3-VSVG-EGFP were incubated at 40°C overnight, then shifted to 32°C for up to 120 min or 19.5°C for up to 60 min. Cycloheximide (150 μg/mL) was added during the incubation at 32°C and 19.5°C to inhibit protein synthesis. The former cells were stained with an anti-GFP antibody without permeabilization to detect EGFP-VSVG that reached the cell surface. Cell observations were conducted using an AxioObserver.Z1 microscope (Carl Zeiss).

### Gene editing by CRISPR/Cas9

We inactivated the *ATP9A* gene in HeLa cells using the CRISPR/Cas9 system (Tanaka et al., 2016). Target sequences for the *ATP9B* gene were designed using the CRISPR Design Tool from the Zhang lab (http://crispr.mit.edu/). Complementary oligonucleotides (5′-caccg<u>cgggagccgaccggcacagc</u>-3′ and 5′-aaac<u>gctgtgccggtcggctcccgc</u>-3′; target sequences are underlined) were synthesized and introduced into the *Bbs*I-digested PX459 vector (Addgene #48139). To edit the *ATP9B* gene, we employed a method previously described (Nozaki et al., 2017; Tanaka et al., 2016), which involved co-transfecting the plasmid containing *ATP9B* target sequences and the Cas9 gene, and a donor plasmid (pDonor-tBFP-NLS-Neo; deposited in Addgene, # 80766), referred to as Donor (Supplementary Figures S2 and S4). Both plasmids were introduced into HeLa cells via transfection using the X-tremeGENE9 DNA Transfection Reagent (Roche). Transfected cells were selected in a medium containing G418 (1 mg/mL), and clones were isolated based on the expression of the reporter gene Tag-BFP using the SH800S cell sorter. To confirm *ATP9B* editing, genomic DNA was extracted from individual clones and subjected to PCR to amplify the region of interest. The KO was confirmed by direct sequencing of the amplified PCR product, revealing mutations in two alleles (Supplementary Figure S2). The joint sequence between the Donor and *ATP9B* gene was confirmed in clone 1, but not in clone 4, despite the Donor being inserted into the *ATP9B* gene area of chromosome 18 (Supplementary Figure S2). Clone 1 was used for further rescue experiments in this study. For *ATP9A/ATP9B*-DKO, *ATP9A*-KO (clone 23) cells were transfected with the two plasmids as described above, cells were selected in a medium containing puromycin (1 μg/mL), and clones were isolated using the SH800S cell sorter. To confirm *ATP9B* editing, genomic DNA was extracted from individual clones and subjected to PCR to amplify the region of interest. The KO was confirmed by direct sequencing of the amplified PCR product, revealing mutations in two alleles (Supplementary Figure S4).

### Immunoblot analysis

Cells were lysed in a lysis buffer (20 mM HEPES, pH 7.4, 150 mM NaCl, 1% Nonidet P-40, 1 mM EDTA) with a protease inhibitor mixture (Nacalai Tesque) at 4°C for 60 min. The lysates underwent centrifugation at maximum speed for 20 min at 4°C in a microcentrifuge to remove cellular debris and insoluble materials. Cell lysates (10 µg/sample) were incubated in SDS sample buffer containing β-mercaptoethanol at 37°C for 2 h or at RT overnight. Subsequently, the samples were subjected to SDS-PAGE and immunoblot analysis using rat anti-HA, mouse anti-actin, or mouse anti-TfnR antibodies. Immunoblots were developed using ImmunoStar reagents (Wako) and recorded on an ImageQuant 800 instrument (Amersham).

### Blue Native PAGE (BN-PAGE)

A NativePAGE^TM^ Bis-Tris Gel system (Thermo Fisher Scientific) was utilized for BN-PAGE (Wittig et al., 2006) (Suzuki et al., 2016). Cells were lysed with a lysis buffer [20 mM Tri-HCl (pH 7.5), 50 mM KCl, 1 mM MgCl_2_, 1% glycerol, 1% n-dodecyl-β-D-maltopyranoside (DDM), protease inhibitor cocktail (Nacalai Tesque)] on ice for 1 h. Cell lysates (∼15 μg) were loaded onto NativePAGE^TM^ 4 to 16% Bis-Tris Gels. After electrophoresis at 150 V at 4°C for 30 min, the concentration of CBB G-250 in the cathode running buffer was reduced from 0.02% to 0.002%, and the sample was further electrophoresed at 150 V for 90 min. Protein standards (Thermo Fisher Scientific) were used as molecular weight markers. The gels were soaked in SDS-PAGE running buffer (25 mM Tris, 192 mM glycine, and 0.1% SDS) at room temperature for 20 min and then transferred to PVDF membranes (Immobilon-P, Millipore). Membranes were probed with anti-HA or anti-DYKDDDDK antibodies, followed by incubation with HRP-conjugated anti-rat or anti-mouse antibodies. Immunoblots were developed using ImmunoStar reagents (Wako) and recorded on an ImageQuant 800 instrument (Amersham).

## ACKNOWLEDGMENTS

We thank Dr. Jun Suzuki and Ms. Niu Han (Kyoto University) for their technical support with BN-PAGE, and Dr. Toshio Kitamura (The University of Tokyo), Dr. Hiroyuki Miyoshi (RIKEN BioResource Center), and Dr. Peter McPherson (McGill University) for providing materials. This work was supported by JSPS KAKENHI Grant Numbers JP23H02434 and JP23K27127 (to H.-W.S.), the Takeda Science Foundation (to H.-W.S.), the Mizutani Glycoscience Foundation (to H.-W.S.), and the ONO Medical Research Foundation (to H.-W.S.).

## Abbreviations

P4-ATPase: type IV P-type ATPase
PS: phosphatidylserine
PE: phosphatidylethanolamine
PC: phosphatidylcholine
SM: sphingomyelin
VSVG: vesicular stomatitis virus glycoprotein
TGN: *trans*-Golgi network

## Supplemental Figure Legends

**Supplementary Figure S1.**
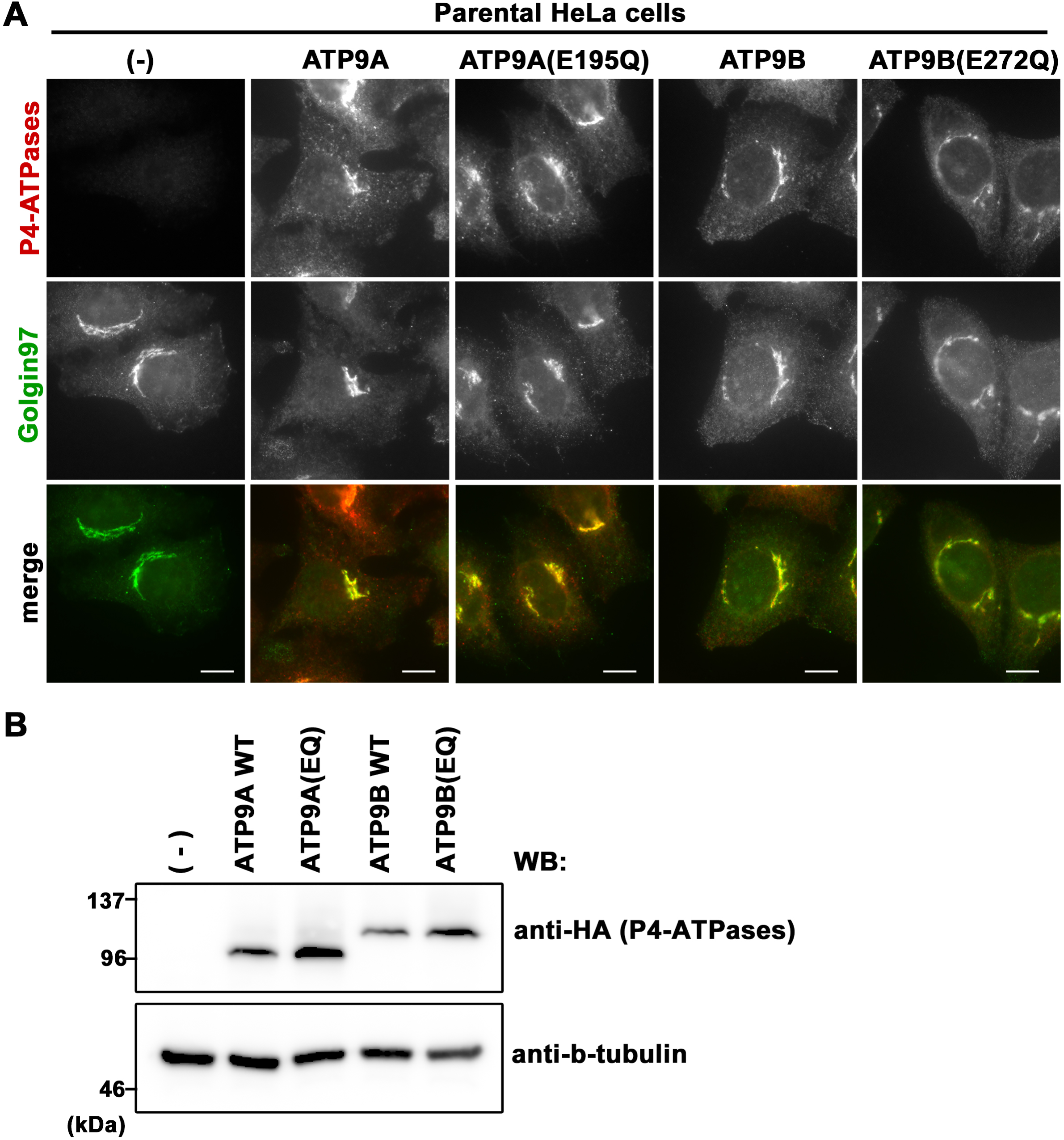
Expression and subcellular localization of ATP9A, ATP9B, and their ATPase-deficient E-to-Q mutants. (A) Parental HeLa cells stably expressing C-terminally HA-tagged ATP9A, ATP9A(E195Q), ATP9B, and ATP9B(E272Q) were fixed, permeabilized, and incubated with anti-HA and anti-Golgin97 antibodies, followed by incubation with Cy3-conjugated anti-rat and AlexaFluor 488-conjugated anti-mouse secondary antibodies. Bars, 10 µm. (B) The expression levels of each ATP9 protein were analyzed by immunoblotting with anti-HA and anti-β-tubulin antibodies.

**Supplementary Figure S2.**
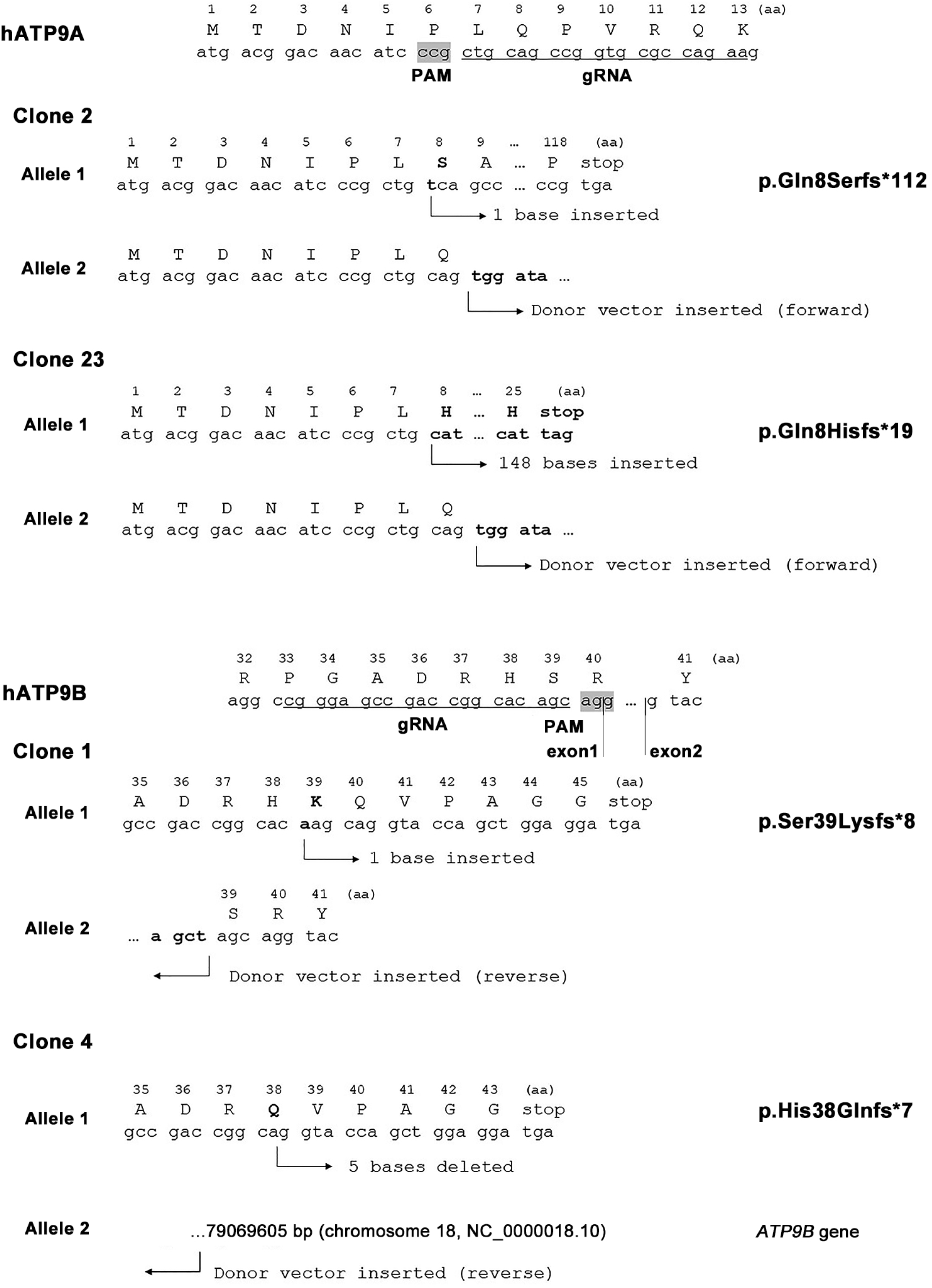
Disruption of *ATP9A* or *ATP9B* by CRISPR/Cas9 in HeLa cells. Two clones of *ATP9A*-KO (clones 2 and 23) were previously described (Tanaka et al., 2016). To confirm the editing of *ATP9B* in HeLa cells, genomic DNA was extracted from individual clones and subjected to PCR. The KO was confirmed by direct sequencing of the amplified PCR products. *ATP9B*-KO (clone 1) carries a biallelic edition with one indel and the insertion of the Donor. *ATP9B*-KO (clone 4) carries a biallelic edition, one indel, and the insertion of the Donor into chromosome 18, where *ATP9B* is located. While the specific joint sequences between *ATP9B* and the Donor were not characterized, it is likely that the Donor inserted upstream of 79,069,605 base pairs (chromosome 18, NC_0000018.10). *ATP9A*-KO (clone 23) and *ATP9B*-KO (clone 1) were used for further rescue experiments.

**Supplementary Figure S3.**
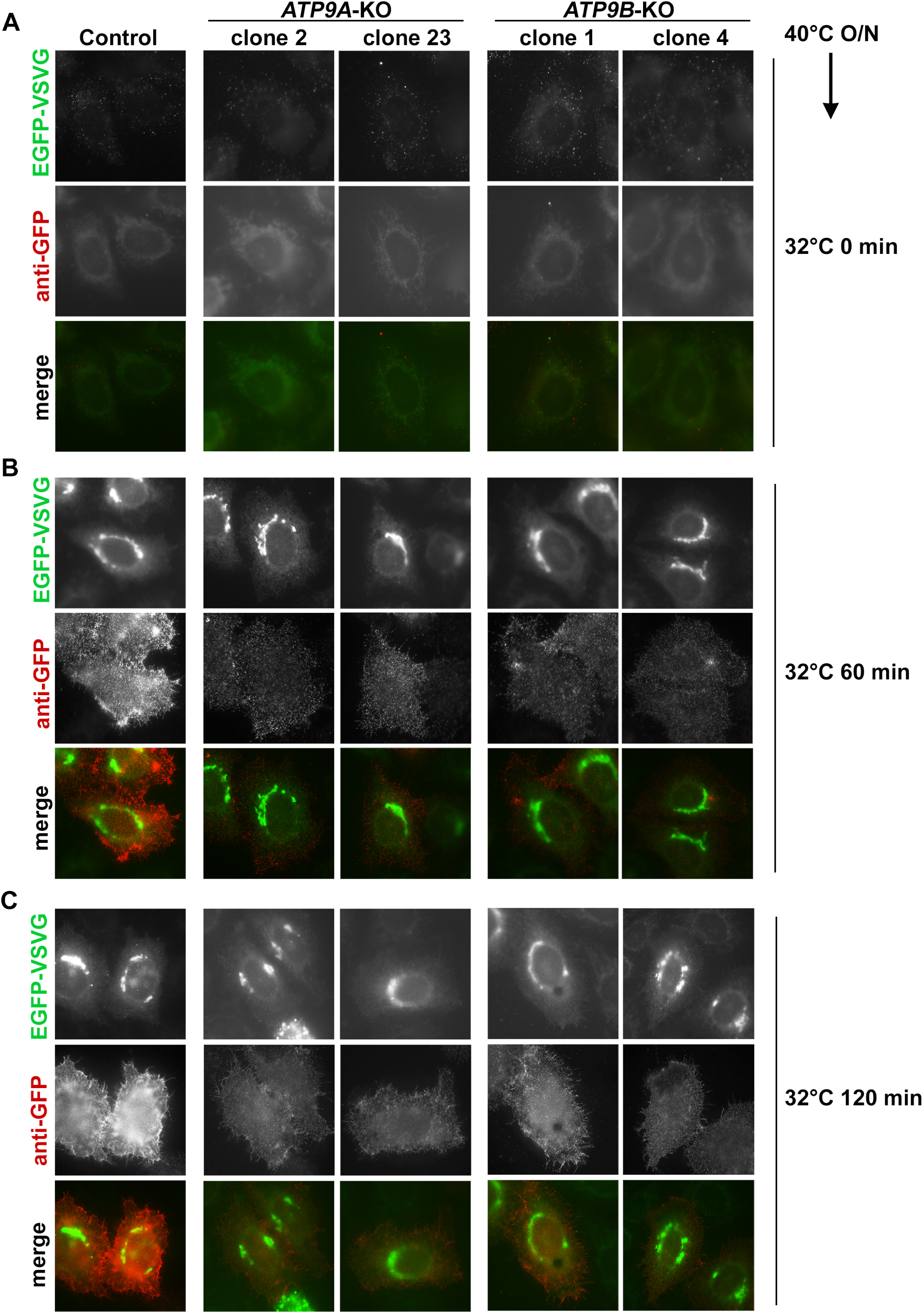
Delay in VSVG(*ts*O45) transport to the plasma membrane in *ATP9A*-and *ATP9B*-KO cells. Parental HeLa (Control), along with two distinct *ATP9A*-KO and *ATP9B*-KO cell clones, were transfected with an expression vector for N-terminally EGFP-tagged VSVG(*ts*O45). Following this, the cells were incubated at 40°C overnight to accumulate EGFP-VSVG at the ER, then shifted to 32°C for the indicated times (**A–C**). Cell surface expression of EGFP-VSVG was detected by staining the cells with anti-GFP antibody under non-permeabilized conditions, followed by AlexaFluor 555-conjugated anti-rabbit secondary antibody (red). N-terminally EGFP-tagged VSVG was barely detectable in the ER after 40℃ incubation due to EGFP misfolding (**A**), but EGFP-VSVG became visible after 32℃ incubation (**B, C**). Bars, 10 µm.

**Supplementary Figure S4.**
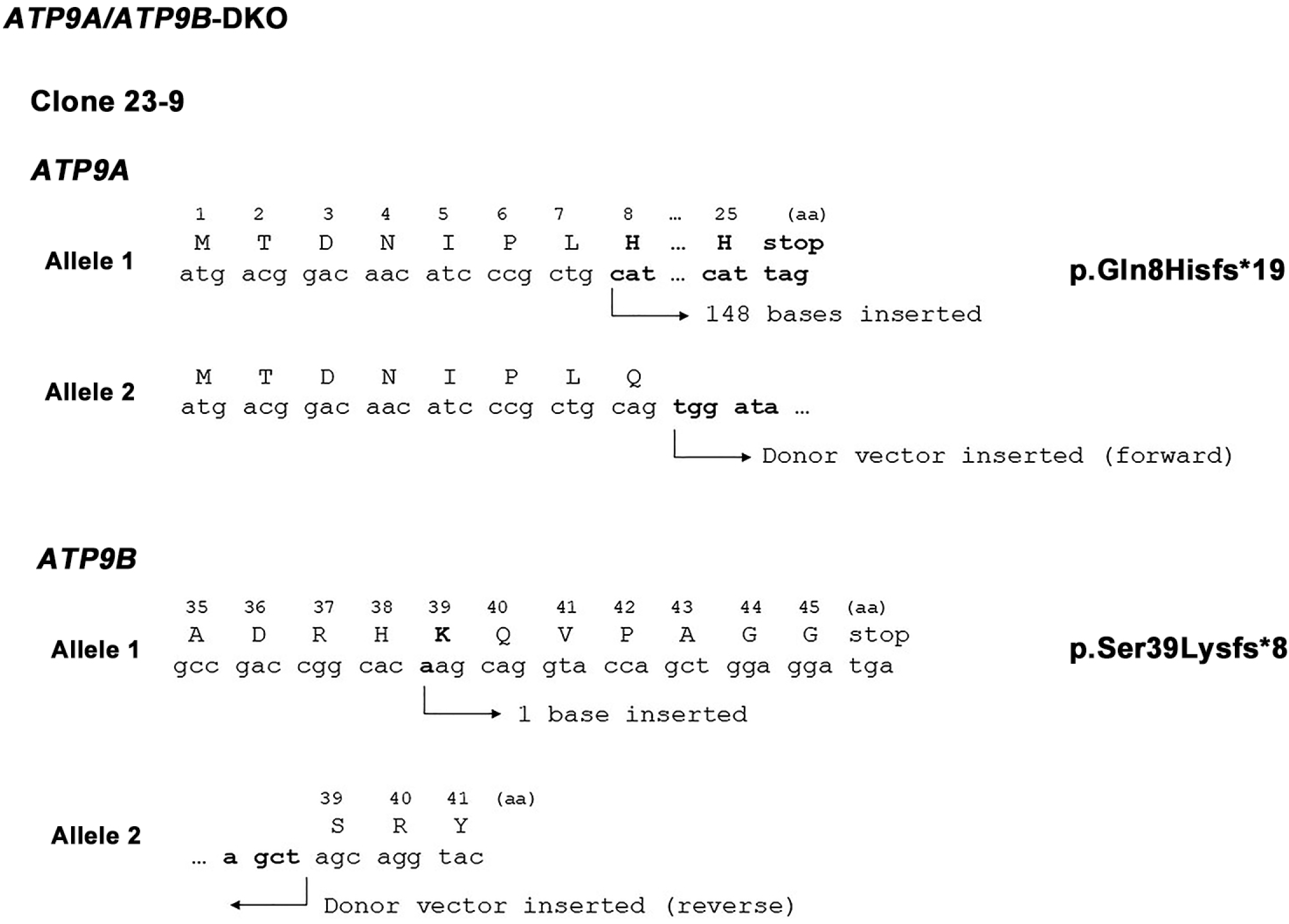
*ATP9A* and *ATP9B* disruption by CRISPR/Cas9 in HeLa cells. To generate *ATP9A/ATP9B*-DKO cells, *ATP9B* was disrupted in *ATP9A*-KO (clone 23) cells. To verify the editing of *ATP9A* and *ATP9B* in HeLa cells, genomic DNA was extracted from individual clones and subjected to PCR. The KO was confirmed by direct sequencing of the amplified PCR products. In the *ATP9A/ATP9B*-DKO (clone 23-9) cells, *ATP9B* carries a biallelic edition, an indel, and the Donor, as well as *ATP9A,* as shown in Supplementary Figure S2.

**Supplementary Figure S5.**
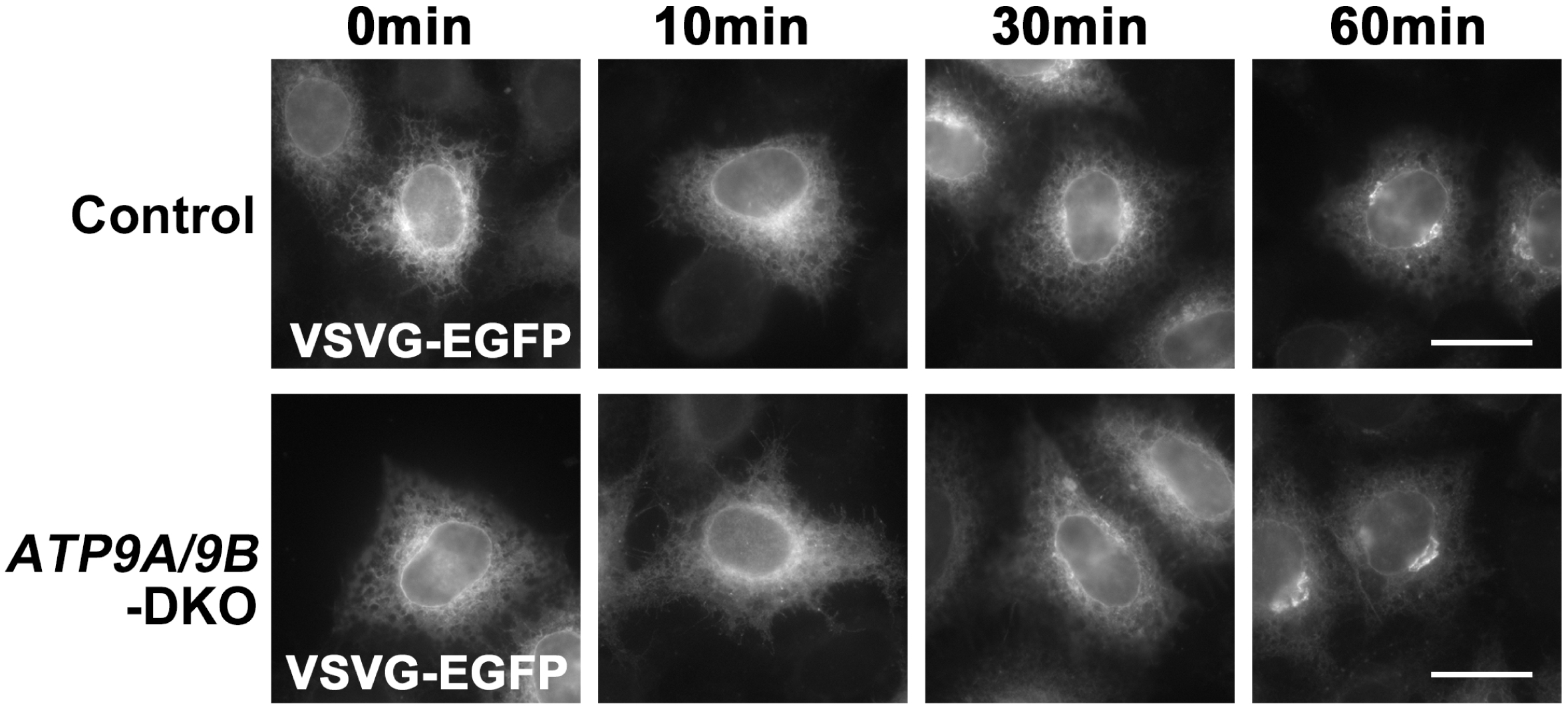
ER-to-Golgi transport of VSVG(*ts*O45) remains unaffected in *ATP9A*/*ATP9B*-DKO cells. Parental HeLa (Control) and *ATP9A*/*ATP9B*-DKO cells were transfected with an expression vector for C-terminally EGFP-tagged VSVG(*ts*O45). Following this, the cells were incubated at 40°C overnight and shifted to 19.5°C for the indicated times. VSVG-EGFP is detectable in the ER after 40°C incubation and appears in the Golgi area upon 19.5°C incubation. Cycloheximide was administered during the 19.5°C incubation. Bars, 20 µm.

**Supplementary Figure S6.**
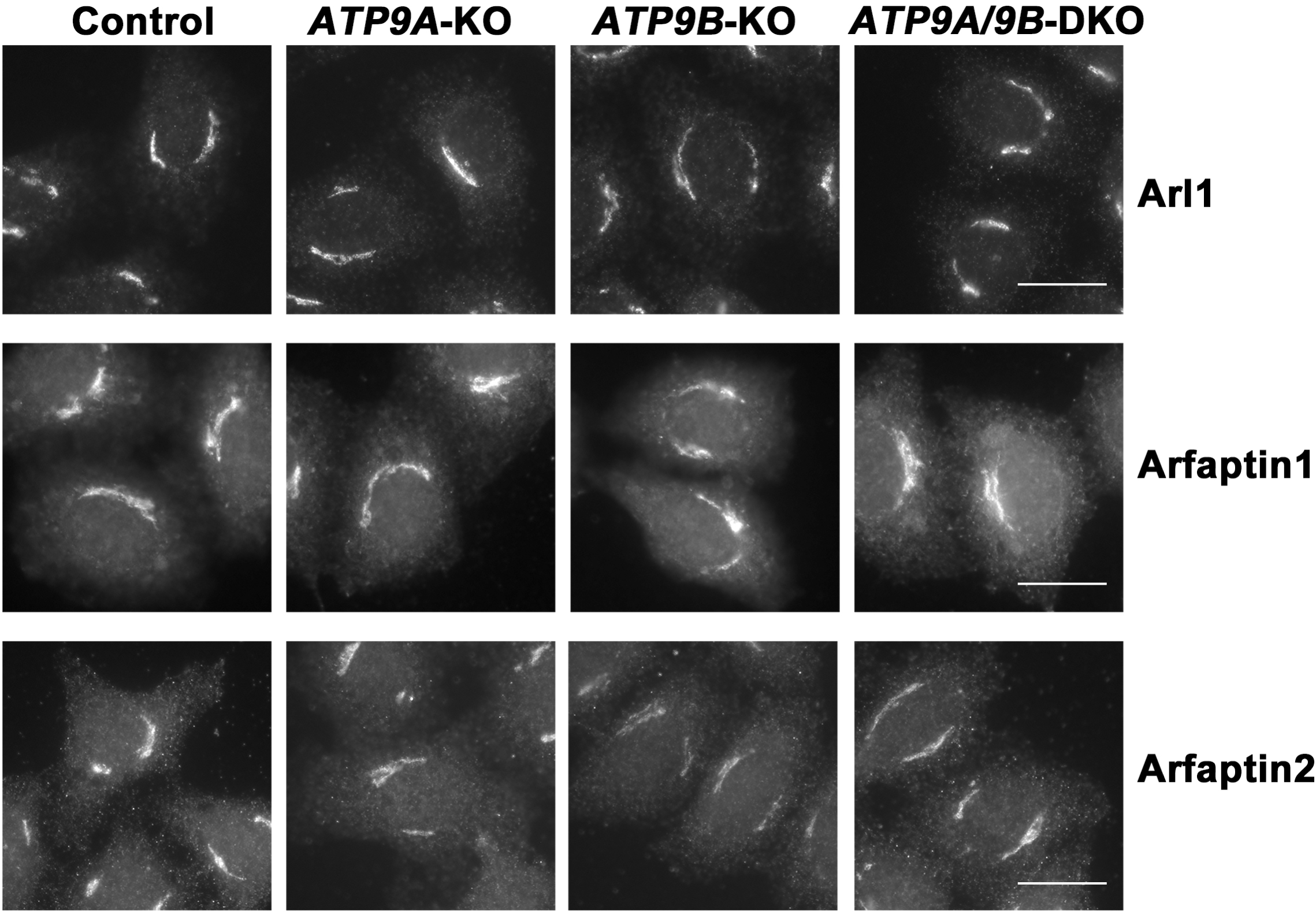
Golgi-localizing proteins recruitment is largely unaffected in ATP9A and/or ATP9B depleted cells. The parental HeLa (Control), *ATP9A*-KO, *ATP9B*-KO, and *ATP9A/ATP9B*-DKO cells were fixed, permeabilized, and incubated with antibodies against Arl1, Arfaptin1, or Arfaptin2. Bars, 20 µm.

